# Ultra-high throughput mapping of genetic design space

**DOI:** 10.1101/2023.03.16.532704

**Authors:** Kshitij Rai, Ronan W. O’Connell, Trenton C. Piepergerdes, Yiduo Wang, Lucas B. C. Brown, Kian D. Samra, Jack A. Wilson, Shujian Lin, Thomas H. Zhang, Eduardo M. Ramos, Andrew Sun, Bryce Kille, Kristen D. Curry, Jason W. Rocks, Todd J. Treangen, Pankaj Mehta, Caleb J. Bashor

## Abstract

Massively parallel genetic screens have been used to map sequence-to-function relationships for a variety of genetic elements. However, because these approaches only interrogate short sequences, it remains challenging to perform high throughput (HT) assays on constructs containing combinations of multiple sequence elements arranged across multi-kb length scales. Overcoming this barrier could accelerate synthetic biology; by screening diverse gene circuit designs and learning “composition-to-function” mappings that reveal genetic part composability rules and enable rapid identification of behavior-optimized variants. Here, we introduce CLASSIC, a genetic screening platform that combines long- and short-read next-generation sequencing (NGS) modalities to quantitatively assess pools of constructs of arbitrary length containing diverse part compositions. We show that CLAS-SIC can measure expression profiles of >10^5^ gene circuit designs (from 5-20 kb) in a single experiment in human cells. The resulting datasets can be used to train ML models that accurately predict circuit behavior across expansive circuit design landscapes, revealing part composability rules that govern circuit performance. Our work shows that by expanding the throughput of each design-build-test-learn (DBTL) cycle, CLASSIC enhances the pace and scale of synthetic biology and establishes an experimental basis for data-driven design of complex genetic systems.

## MAIN TEXT

Synthetic gene circuits are constructed by assembling DNA-encoded genetic parts into multi-gene programs that perform computational tasks in living cells^1–3^. Over the past two decades, gene circuits have emerged as important models for understanding native gene regulation^4^ and have been used to create powerful biotechnologies by enabling user-defined control over cellular behavior^5–7^. Despite this progress, the design of quantitatively precise circuit behavior remains challenging. Regulatory interactions within a circuit must be carefully tuned, often through multiple iterative DBTL cycles, before part compositions that support a desired circuit behavior are identified^8^. Additionally, since genetic parts must work in close physical proximity to one another, as well as within a crowded intracellular environment^9^, incidental molecular coupling can occur between parts and with host cell regulatory machinery^10–12^. These context-dependent interactions can confound the ability to design circuits using biophysical and mechanistic modeling frameworks, further extending the number of DBTL cycles required to achieve a target behavior^13^. One potential strategy for increasing the pace of gene circuit engineering is to expand the number of circuits tested in each cycle by performing HT functional screens on pooled circuit libraries. By measuring large collections of circuits in a single experiment, such an approach could be used for rapidly profiling complex circuit design spaces to identify part compositions with desired quantitative behaviors. In contrast to traditional circuit engineering approaches, collecting large amounts of circuit activity data could also facilitate the training of ML/AI models that are capable of inferring context-specific part function and more accurately predicting the behavior of untested circuit designs. HT screening approaches that utilize short-read NGS as a readout^14–18^ have been used to generate detailed sequence-to-function mappings for multiple genetic part classes, including promoters^19–23^, terminators and 3’ UTRs^24,25^, transcription factors (TFs)^26,27^, nucleic acid switches^28^, and receptors^29^. However, high-depth functional profiling of libraries of DNA constructs long enough (>1 kb) to encode entire circuits can be costly and technically challenging. Methods have emerged that permit multiplexed analysis of long constructs by appending short barcode index sequences that facilitate amplicon-based readout by short-read NGS^30^. This approach has been used in array formats in which each construct is assembled and barcoded separately, or through nested assembly schemes^30,31^, limiting library design flexibility and assay throughput. More recently, random barcoding and indexing via long-read single-molecule realtime (SMRT) sequencing (PacBio) has been used to characterize libraries of variants ranging from 10^2^ to 10^4^ constructs, mostly for individual ORFs of up to 6 kb in length^32–34^.

To perform indexed multiplexing for construct libraries containing combinations of genetic parts at larger length scales, we devised an approach that combines long-read nanopore and short-read Illumina NGS to analyze construct libraries generated via pooled part assembly (**Fig. 1A**). Our assembly scheme incorporates structured barcodes that can be used to create a construct composition-to-barcode index by nanopore sequencing. The construct libraries can then be introduced into cells, binned based on functional phenotype, and analyzed by short-read (Illumina) amplicon sequencing to produce a barcode-to-phenotype index. A map that matches construct composition to phenotype can be made by comparing the two indices. Using this technique, which we refer to as CLASSIC (<u>c</u>ombining long-<u>a</u>nd <u>s</u>hort-range <u>s</u>equencing to investigate genetic <u>c</u>omplexity), it is possible to obtain high-depth phenotypic expression data for large libraries (>10^5^) of genetic part compositions of arbitrary length using standard phenotypic selection or flow sorting experiments.

**Figure 1.**
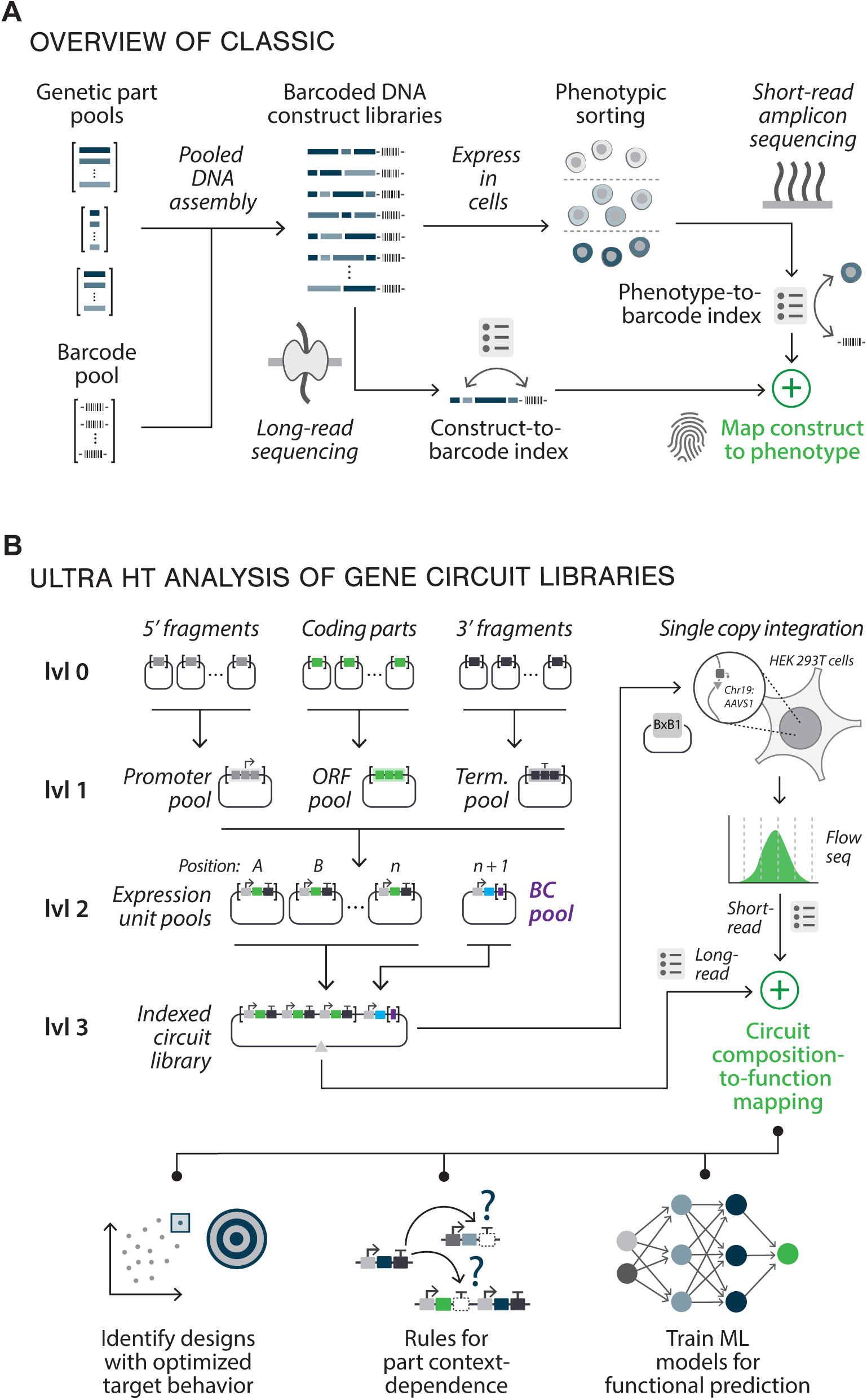
Using CLASSIC to systematically map the design space of complex genetic programs. (**A**) Overview of CLASSIC. Pooled assembly of genetic parts with DNA barcode sequences yields libraries of barcode-indexed constructs of arbitrary length and complexity. Long-read nanopore sequencing is used to create an index matching construct composition to an associated barcode. In parallel, libraries introduced into cells undergo sorting or selection to bin expression phenotypes. Barcode amplicons generated for each bin are subjected to short-read NGS to quantify expression phenotype, which is then mapped to construct composition via barcode indexes. (**B**) Application of CLASSIC to profile a synthetic gene circuit design space. Hierarchical golden gate assembly is used to compose libraries of multi-EU circuits with combinatorially varied part compositions and circuit designs. Sequence fragment pools for different parts categories from level 0 are combined to yield level 1 pools of promoters, open reading frames (ORF), and terminators (Term.), which are then combined to yield level 2 EUs (square brackets: fragment/part pools). Barcode (BC) pools harbored on BFP EUs are combined with library EU pools to create indexed multi-EU circuit libraries (level 3) that are integrated into HEK293T-LP cells placed at the *AAVS1* locus in chromosome 19 via expression of the BxB1 recombinase (top right). Library analysis by a combination of nanopore and flow-seq yields composition-to-function mapping.

To configure CLASSIC to quantitatively profile libraries of constructs containing diverse genetic part combinations in human cells (**Fig. 1B**), we designed a custom hierarchical golden-gate^35^ cloning scheme in which library diversity is programmed through a series of pooled DNA assembly steps. Diversified pools of 5’, coding, and 3’ gene elements are first generated through the assembly of input part fragments (level 0 to 1), and subsequently combined to create single-gene expression unit (EU) pools (level 1 to 2). The EU pools are then combined with a BFP EU plasmid pool that expresses a transcript containing semi-degenerate barcode sequences in the 3’ UTR, yielding barcoded multi-EU circuit libraries (level 2 to 3). Following nanopore sequencing and data analysis (see **Methods**), the libraries are genomically integrated at single copy into a HEK293T cell line harboring a custom “landing pad” cassette (HEK-LP)^36,37^ (see **Methods**). Library-integrated cells are then flow-sorted based on circuit expression output, followed by Illumina NGS to quantitate bin distributions of circuit-associated barcodes^14,15^ (see **Methods**). We validated this workflow with a small (n=384) expression unit (EU) library containing shuffled regulatory elements (promoter^38,39^, Kozak sequence^40^, and terminator^24,41,42^), demonstrating that our assembly scheme yields complete, part-balanced library coverage and proportional assortment of unique barcodes. HEK-LP library integration and flow-seq measurements were compared with flow cytometry measurements made with a set of randomly isolated, clonally expanded variants. These “ground truth” measurements showed excellent agreement with corresponding CLASSIC-derived values (mean absolute error [*MAE*]=0.07), and demonstrated that measurement precision is enhanced by mapping multiple unique barcodes to each composition. We also collected replicate libraries to demonstrate that CLASSIC is technically (*r^2^*=0.99) and biologically (*r^2^*=0.97) reproducible. Finally, we showed that the data can be used to quantify relative part contributions to expression level, as well as identify unexpected interference between parts.

To evaluate whether CLASSIC can be used to experimentally profile a complex design space for a multi-gene circuit, we constructed a library of small-molecule inducible, single-input transcriptional switches comprising >10^5^ potential unique compositions. Circuits in this library consist of two EUs (**Fig. 2A**): one encoding a constitutively expressed synthetic Cys2-His2 zinc-finger (ZF)-based transcription factor (synTF)^43–45^ with appended transcriptional activation domains (TAs), and the other encoding a reporter gene harboring cognate synTF binding motifs (BMs) located upstream of a minimal promoter driving eGFP expression^43^. Transcriptional induction occurs upon addition of the small molecule 4-hydroxytamoxifen (4-OHT), which binds to a mutant version of the human estrogen receptor (ERT2)^46^ appended to the synTF, facilitating its translocation from the cytoplasm into the nucleus to activate reporter transcription. As we and others have demonstrated^43,47–49^, identifying designs that are optimized for high fold-change (HFC) (**Fig. 2A, right**) expression can be challenging in mammalian systems; to ensure robust expression of a transgene exclusively in the presence of inducer, basal and induced expression levels must be respectively minimized and maximized through tuning of both the transcriptional regulatory features of the locus and the molecular properties of the synTF.

**Figure 2.**
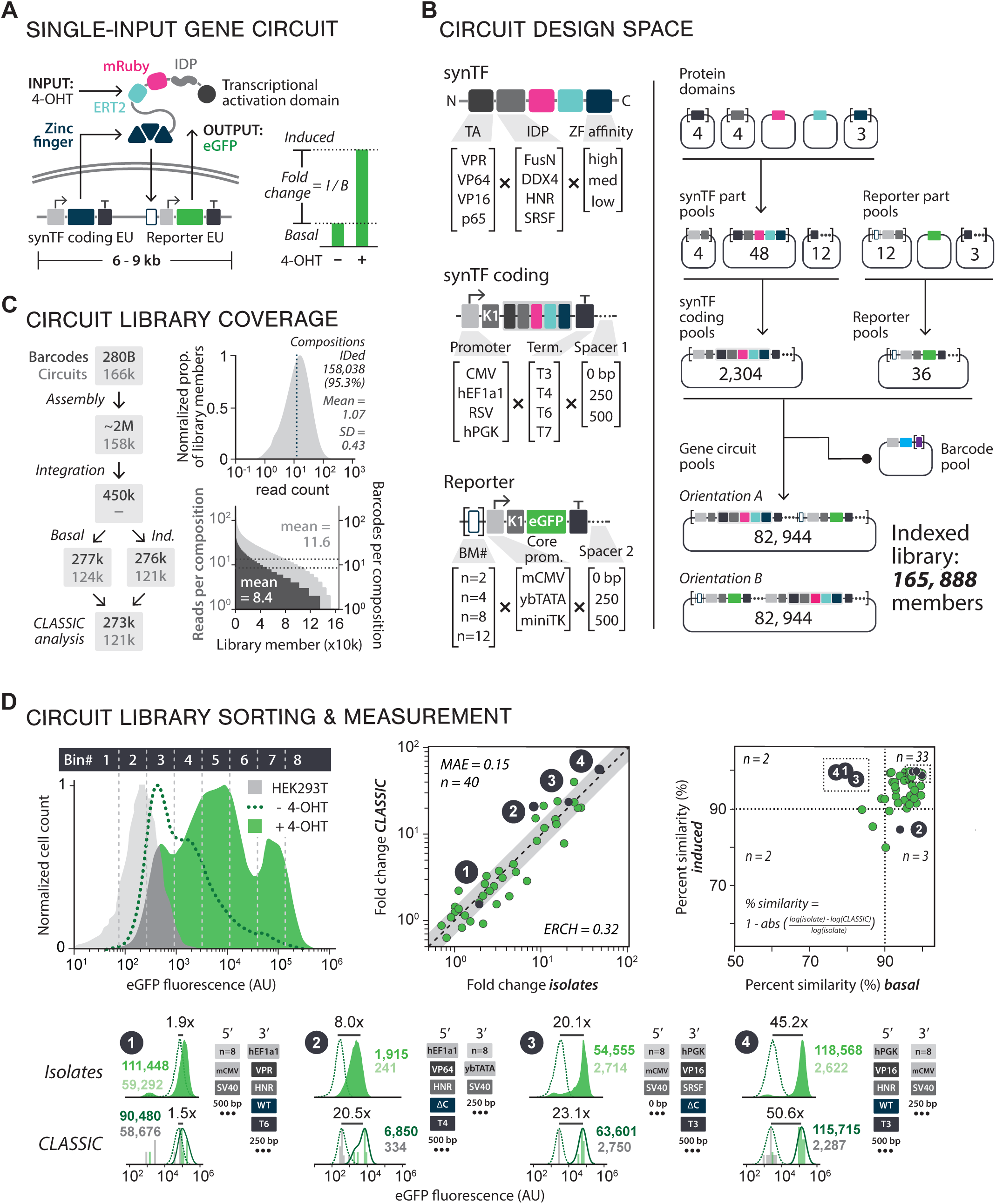
Using CLASSIC to quantitatively profile a synthetic gene circuit design landscape. (**A**) Inducible synTF circuit diagram. Left: The circuit contains two EUs: one codes for the synTF and the other is an eGFP reporter. Without inducer, the synTF localizes to the cytoplasm. Upon addition of 4-OHT (input), the synTF translocates into the nucleus to bind to and activate reporter eGFP expression (output). Right: Expression fold change is the ratio of expression levels in the presence or absence of inducer. (**B**) synTF circuit design space. Left: SynTF diversity (arranged N-to-C term): of 4 TAs, 4 IDPs, and 3 ZF affinity mutants (high: WT, med: ΔC, low: ΔC 6x). synTF coding EU diversity: 4 constitutive promoters, 4 terminators. Reporter EU diversity: 4 BM number variants, 3 minimal core promoters. This unit contains a constant terminator. Both EUs have 3’ spacing sequences of 0bp, 250bp or 500bp downstream. Right: EUs assembled from input parts and combined in 2 different 5’-to-3’ orientations, to generate a combinatorial diversity of 165,888 possible circuit variants. (**C**) Balance of circuit library assembly and indexing. Left: data for ∼121k out of 166k variants (73%) were recovered by CLASSIC; compositions, light grey; barcodes, dark grey. Right: Nanopore sequencing data of the library were analyzed to assess library composition count (mean and standard deviation [SD] computed on log_10_ scale) (top), composition/barcode balance (reads per composition [light grey], unique barcodes per composition [dark grey]; data plotted in rank order of reads per composition) (bottom). (**D**) Circuit library sorting and measurement. Top right: eGFP expression of circuit library in presence (solid green) and absence (dotted green line) of 4-OHT (top, center) is shown, along with boundaries of flow sorting bins (vertical grey dotted lines); grey histogram, empty HEK293T-LP cells. Top middle: Fold-change values for 40 clonal isolates plotted against CLASSIC-derived values. Black dots, isolates displayed at the bottom of the panel; *MAE*, mean average error; grey region, ERCH, error range of clonal heterogeneity AU, arbitrary fluorescence units. Top right: % similarity between basal and induced values for CLASSIC and isolates. Bottom: Flow data from 4 isolates (green dotted line, uninduced; green solid, induced) are shown with corresponding CLASSIC-computed distributions [kernel density (black line) calculated from normalized barcode read count (grey vertical bars)]. Parts combinations corresponding to each index are shown.

To create a library that diversifies the design of this circuit architecture, we identified 10 genetic part categories that we hypothesized could play a role in circuit tuning to achieve HFC behavior. This included 4 different transcriptional activation domains (TAs)^50–52^, 3 ZF affinities^43^, and a set of 4 intrinsically disordered protein (IDP) domains^53–55^ that have been shown to facilitate liquid-liquid phase condensation of nuclear-localized TFs^53,54^ (**Fig. 2B, left**). We also varied transcriptional regulatory features of the circuit, including 4 promoters and 4 terminators in the synTF coding EU, as well as 3 core promoters^56–58^ and different numbers of BMs (2, 4, 8, or 12) in the reporter EU (**Fig. 2B, left**). Additionally, we varied the spacing between the EUs (0, 250, or 500 bp) along with their 5’-to-3’ orientation (**Fig. 2B, left**) to yield an overall circuit design space of 165,888 compositions (**Fig. 2B, right**). This library was constructed in 3 steps by first assembling protein domain parts to create a level 1 synTF ORF pool, then conducting parallel assemblies to generate level 2 pools of synTF coding and reporter EUs, and finally combining EU and barcode pools into the level 3 destination vector (**Fig. 2B right**).

Nanopore sequencing of the pooled library yielded barcode assignments for 95.3% of total compositions (**Fig. 2C**), with a mean of 8.4 barcodes for each circuit (**Fig. 2C bottom left**). We integrated the library into HEK-LP cells and sorted un-induced and 4-OHT-induced populations (8.6 and 15.7 million cells, respectively) separately into 8 bins based on eGFP fluorescence (**Fig. 2D left**). Following analysis by Illumina NGS, we matched a total of 121,292 (73% of design space) compositions to barcodes that were detected in the sorted populations (mean barcodes circuit^-1^=2.25) (**Fig. 2C**). We then used barcode bin distributions to compute basal, induced, and fold-change expression values for each variant. CLASSIC-derived fold-change values demonstrated excellent overall agreement (*MAE*=0.15) (**Fig. 2D top middle**) with those of random isolates (n=40) (**Fig. 2D bottom**), and values for both basal and induced expression showed comparably high percent similarities (**Fig. 2D top right**). These results confirm that CLASSIC can generate quantitatively accurate measurements of libraries large enough to profile complex genetic design spaces.

Since an incomplete mapping of circuit design space (27% was unmeasured) could limit our ability to identify HFC-optimized compositions, we tested whether a complete mapping could be obtained by training an ML model with our data and then using it to predict the behavior of unmeasured compositions. We trained several model classes and found that multi-layer perceptron (MLP) (90/10 train/validation split, with high-quality test set *r^2^*=0.86 for basal 0.90 for induced data) and random forest (0.84 and 0.86) models significantly out-performed biophysically-consistent (0.46 and 0.27) and unconstrained linear regression models (0.53 and 0.64), indicating that the complex, non-linear interactions between genetic parts can be more accurately captured by advanced model architectures. We selected the MLP to move forward with (**Fig. 3A**) due to its predictive accuracy and potential for extensibility (**Fig. 3A)** (see **Methods**). We used the MLP to impute basal and induced eGFP expression values to yield a complete design space mapping of all 165,888 compositions (**Fig. 3B**). We then analyzed the distribution of values across a 2-dimensional projection of behavior space (basal vs. induced expression), examining the relative densities of compositions in 3 regions of interest: low basal (<500 AU), high induced (>70,000 AU), and HFC expression (>25x fold-change) (**Fig. 3B, right**). We observed a greater proportion of library members in the low basal than in the high induced region (∼4.3x), with ∼50% of compositions showing fold-change values of <3x. Only a small fraction of the library (∼8%) fell within the HFC region. While a comparison between MLP-modeled and CLASSIC-measured distributions revealed similar global features, we observed high absolute error (>2) between the model predictions and CLASSIC-measured values at the periphery of behavior space (∼3% of variants, including many in the HFC region), suggesting either CLASSIC measurement errors or poor model prediction in these regions.

**Figure 3.**
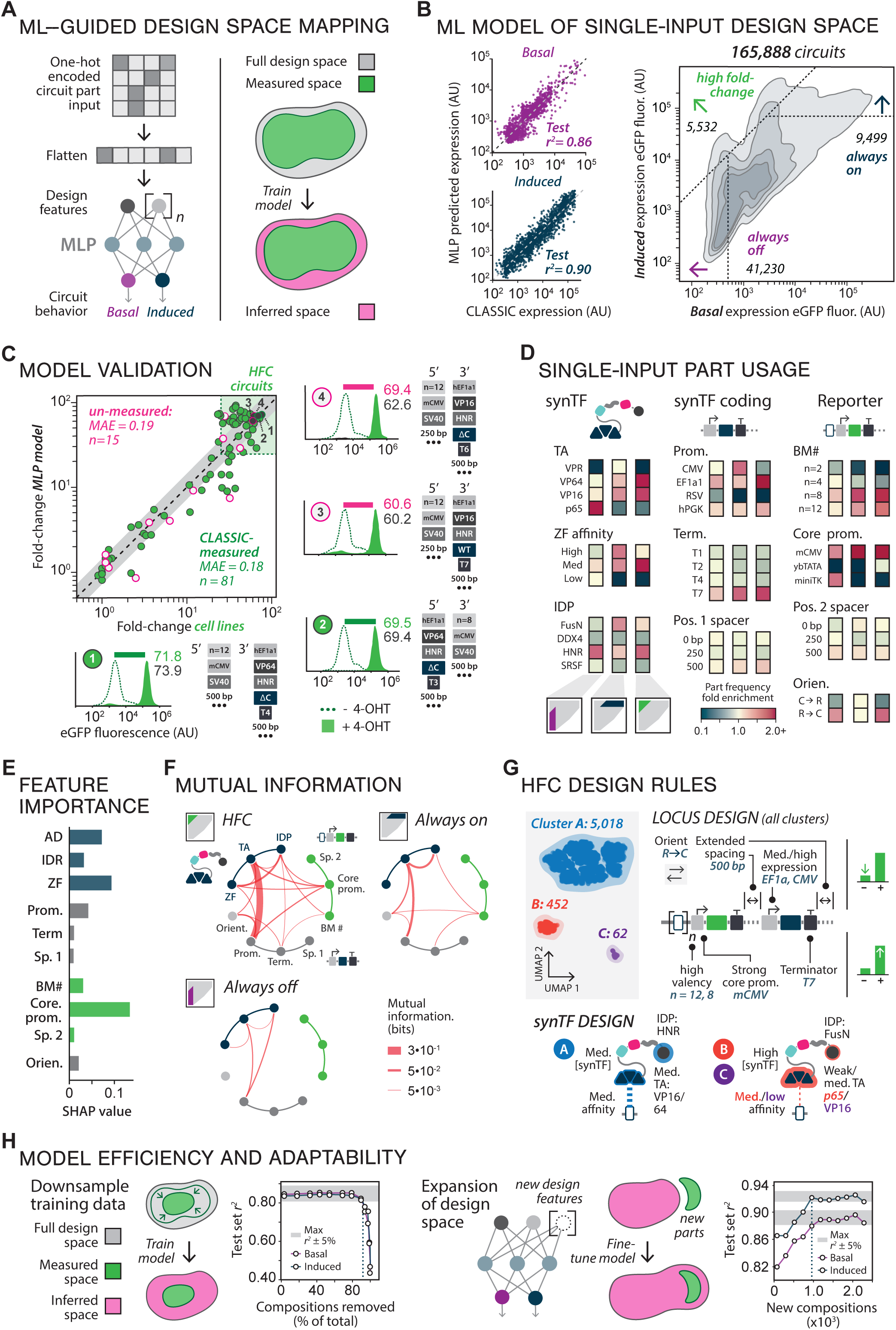
ML-aided mapping of single-input circuit design space reveals gene circuit design rules. (**A**) Left: Multi-layer perceptron (MLP) neural network model of inducible synTF circuit behavior space. Circuit part inputs are one-hot encoded, flattened and passed through a 4-layer MLP with 2 output heads that predict basal and induced expression. Right: ML task description. From the complete design space (gray) of ∼166k compositions, data from 121k was collected (green) and used to train the ML model to re-fit circuit behavior for measured compositions and infer unmeasured space (pink). (**B**) Left: MLP predictions on a held-out high-quality test set (see **Methods**) against basal and induced CLASSIC measurements (purple, basal; blue, induced) (*r*^2^, Pearson’s *r*^2^). Right: Contour plot of MLP modeled basal and induced expression behavior space (contours, light to dark: 97.5%, 90%, 70%, 50% of total measurements). Regions of interest bounded by dotted lines: low basal (<500 AU), purple arrow; high induction (>70k AU) blue; high fold change (HFC) (>25x, green). Values in the plot indicate the number of compositions in each region. (**C**) Experimentally validating ML prediction of circuit function. Top left: Fold-change values for cell lines from classic measured configurations (green filled) and unmeasured configurations (pink outlined) were plotted against CLASSIC-derived values. Black dots, isolates displayed at the periphery of the panel; *MAE*, mean average error; grey region, ERCH, error range of clonal heterogeneity (see **Methods**); green box, HFC behavior region. Right and bottom: Flow cytometry analysis of cell line behavior (green dotted line, basal; solid green, induced) and associated circuit compositions (below each plot). Horizontal pink bars indicate ML predicted fold change. AU, arbitrary fluorescence units. (**D**) Genetic part usage in highlighted regions of behavior space; left to right: low basal, high induced, HFC. Part fold enrichment is calculated by dividing observed part occurrence by expected part occurrence from a balanced library (**E**) Feature importance for all part categories based on absolute SHAP values computed in the HFC region (see **Methods**). Larger SHAP values indicate greater stronger contribution to the model’s predictions of HFC behavior. (**F**) Mutual information between part categories in different regions of behavior space. MI between part categories is denoted by red line thickness. (**G**) Clustering analysis of HFC circuit designs. Part features from HFC designs were subjected to UMAP dimensional reduction followed by K-means clustering with optimal cluster number determined using the gap test (see **Supplementary text**). Top left: UMAP projection of HFC designs. All clusters displayed significant part occurrence similarities in the locus design (top middle), and inter cluster differences in synTF expression level, binding affinity, and activation domain preference (bottom). Right: Mechanism for achieving HFC behavior based on clustering results. Locus features are optimized to reduce basal and maximize induced expression levels, while cluster specific choices in synTF design tend to maximize induced expression. (**H**) Model efficiency and expansion to new features via fine-tuning. Left: The model training set was down-sampled by removing compositions and retraining on smaller percentage of the design space. Middle left: Model performance (Pearson’s *r*^2^) on compositions in the high-quality test set, evaluated on basal (purple line) and induced (green line) expression levels as a function of percentage of total compositions removed from the training set. Dark gray box: maximum observed *r*^2^ ± 5%. Dotted line: number of compositions needed to obtain 95% of maximum observed *r*^2^ value. Middle right: Expansion of the design space by fine-tuning the model on data collected from a library of compositions containing a new part feature (NFZ activation domain). Right: Model performance (test set Pearson’s *r*^2^) for basal and induced expression as a function of increasing number of compositions used in the training set during fine-tuning. Dark gray box: maximum observed *r*^2^ ± 5%. Dotted line: number of compositions needed to obtain 95% of maximum observed *r*^2^ value

To experimentally assess if our model can accurately predict behavior across circuit design space, we constructed cell lines harboring individual CLASSIC-measured (n=81) and unmeasured (n=15) compositions, including many predicted to be among the highest fold-change compositions in the design space (60-80x) (**Fig. 3C**). Flow cytometry analysis of the cell lines closely corresponded with model predictions for both measured (*MAE*=0.18) and unmeasured (*MAE*=0.19) compositions, confirming the behavior of numerous HFC compositions predicted to be in the 60-80x range, near the upper boundary of behavior space. Cell lines constructed for circuit configurations that correspond to apparent high-error CLASSIC measurements also showed strong overall agreement with the model, confirming that it performs accurate error correction for measurement outliers. These results validate the ability of our CLASSIC-trained ML model to enable gene circuit design space mapping through accurate imputation of inaccurately sampled or unsampled compositions.

With a quantitatively accurate mapping of single-input design space in hand, we sought to uncover part composition rules governing circuit behavior. First, we examined relative part usage frequencies in design space regions of interest (**Fig. 3D top**). We observed distinct part usage in categories associated with the synTF protein (TA, IDP, and ZF affinity), as well as the promoter driving synTF expression, the number of BMs, the core promoter, and EU orientation. For example, VPR, a strong TA, is used extensively in the high induced region, but is nearly absent amongst low basal circuits in favor of a weaker TA, p65, while intermediate-strength VP16 and VP64 are enriched amongst HFC circuits. By the same token, lower- and medium-activity core promoters (miniTK and ybTATA) are excluded from high activity circuits, while the low basal region has few compositions with mCMV (high activity core promoter), and HFC circuits utilize a mix of ybTATA and mCMV. HFC circuits favored compositions with the reporter EU in the 5’ position, an orientation that appears to minimize basal circuit expression across design space, potentially by eliminating transcriptional crosstalk from the upstream promoter.

To gain further insight into the design rules underlying HFC behavior, we used our trained MLP to compute Shapley additive explanation (SHAP) values, a widely-used approach that provides an objective metric of each part category’s relative importance in determining target behavior (**Fig. 3E**)^59^. SHAP analysis revealed that HFC behavior is primarily influenced by part selection within six categories: TA, IDP, ZF affinity, and synTF expression and core promoters. To investigate part usage co-variance, we computed cross-category mutual information (MI) for compositions within the HFC region (**Fig. 3F**). This analysis revealed strong coupling among the six SHAP-important categories, indicating that their co-optimization is critical for achieving HFC behavior. To further explore this interdependence, we applied UMAP-assisted K-means clustering^60^ on HFC compositions, identifying three clusters (**Fig. 3G**). Interestingly, all three had distinct synTF designs, but common locus features. This included features that maximize activation like high BM valency (n=8 or 1), a strong core promoter (mCMV), and terminator preference, as well as those that minimize basal expression such reporter-to-coding orientation, medium synTF expression, and extended spacing (500 bp) between EUs (**Fig. 3G**). The largest cluster (A) featured medium-activity promoters driving synTF expression (hEF1a1 and hPGK) and medium-strength TAs (VP64 and VP16) paired with HNR IDP, and medium-affinity ZFs. Performing a dose-response analysis on a representative composition from this cluster revealed a Hill coefficient (*n_H_*) of 2.13, indicating moderate cooperativity. In contrast to cluster A, compositions in clusters B and C favored strong expression (CMV promoter), with B containing medium-affinity ZFs and a weak TA (p65), and C showing enrichment for weak-affinity ZFs and medium-strength TA (VP16). Taken together, these observations suggest that HFC behavior is achieved via a common locus design that minimizes basal reporter expression, and synTF designs that carefully balance expression level, ZF affinity, and transcriptional activation. These results demonstrate that analysis of CLAS-SIC-enabled design space mappings can provide novel insight into how part composition enables target circuit behavior.

While our results establish that accurate design space mappings can be obtained for regimes with balanced training-to-prediction ratios, we realized that models trained with CLASSIC data would offer greater utility if they could generalize to a broader range of out-of-sample test cases. To evaluate the model’s generalization capacity, we progressively downsampled the training data while performing validation with the held-out test set (**Fig. 3H, left**). Surprisingly, we found that only ∼9% of the data was sufficient to achieve an *r^2^*that was 95% of the maximum (>0.81). We then examined whether the design space mapping could be expanded by incorporating additional genetic part measurement data (**Fig. 3H, right**). To do this, we assembled two libraries, each containing approximately 3.5k members: one incorporating a previously unmeasured TA (NFZ)^61^ and another with a 10-GS linker replacing the IDP. Following measurement of these libraries, we fine-tuned the existing “base” model with the new data to predict the expanded design space, which now included the new parts (approximately 259k compositions). While the base model alone showed good predictions of new part behavior (*r^2^*of ∼0.82 for basal and 0.87 for induced conditions for NFZ) the addition of the new data improved model generalization, achieving 95% of maximum accuracy (*r^2^* of ∼0.88 for basal and 0.92 for induced conditions) with just ∼1,000 instances. These results show that our modeling framework is highly data-efficient and can easily and rapidly incorporate new parts and design features through fine-tuning. This suggests its potential, through multiple cycles of library construction and testing, to generalize to design spaces that are much larger than the datasets used for training.

We attempted to evaluate these capabilities for design space expansion by mapping a design space for multi-input circuits with two inducible synTFs, one regulated by 4-OHT and the other by Grazoprevir (GZV), a small molecule protease inhibitor that blocks autolytic cleavage of a NS3 protease-synTF fusion^62^ (**Fig. 4A**). In this 3-EU architecture, co-regulation of the reporter locus by the 2 synTFs could potentially support Boolean logic, including AND-gate function if the synTFs cooperatively activate the reporter, and OR-gate function if they independently maximize activation. We hypothesized that this architecture could be tuned to achieve these behaviors by diversifying a set of 68 design features, including many that were chosen based on our results from the single-input library, as well as synTF BMs arranged in four distinct patterns, yielding in an overall design space of approximately 3.4 billion compositions (**Fig. 4B**). To create a mapping for this space, we employed a two-step data collection strategy, first measuring a densely sampled ”base library” space comprising a 44-feature subset (1.1M compositions), and then obtaining data for an “expansion library” that sparsely sampled the remaining features (**Fig. 4B**). During base library construction, we deliberately constrained diversity to ensure high read depth, obtaining barcode indexes for ∼7% (∼112k compositions) of the 1.1M potential compositions (**Fig. 4C**), with a mean of 3.2 barcodes for each circuit (**Fig. 4C bottom left**). The expansion library was constructed by assembling 24 independent 1,500-member pools, each containing one of the expansion features held constant against a random sampling of all other base library features. Both libraries were integrated separately into HEK-LP cells and treated with 4-OHT, GZV, or both inducers. Each treatment was sorted into 8 bins based on eGFP fluorescence and isolates were obtained from the base (n=14) and expansion libraries (n=13). As before, we performed Illumina NGS to obtain expression values from both libraries (∼83k and ∼45k respectively for base and expansion libraries) (**Fig. 4C, bottom**), once again observing overall agreement between CLAS-SIC and isolate measurements (*MAE*=0.28, 0.23) (**Fig. 4D**).

**Figure 4.**
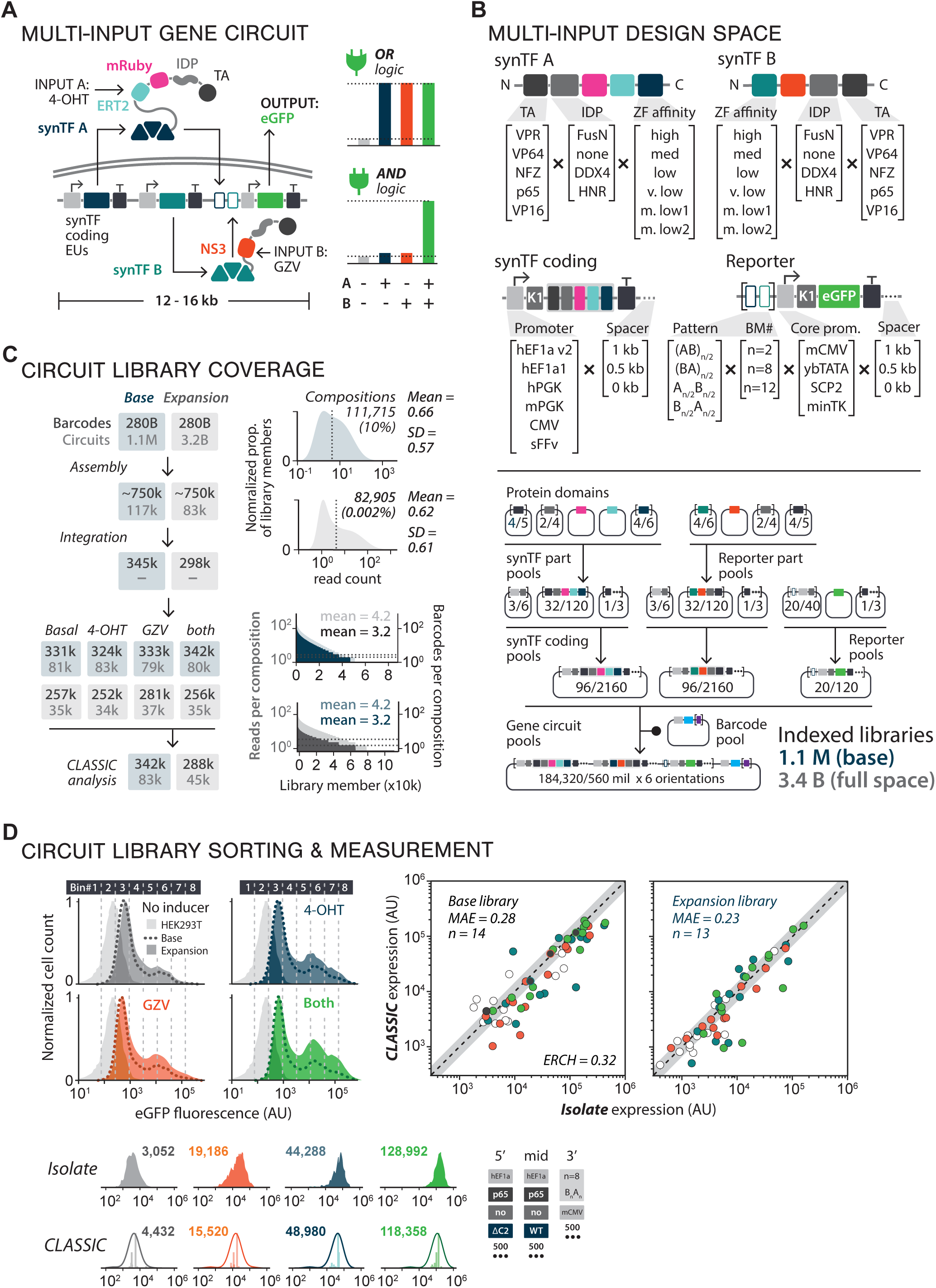
ML guided exploration of multi-input gene circuit behavior. (**A**) Multi-input inducible synTF circuit diagram. Left: the circuit contains three EUs: a 4-OHT inducible synTF A (navy), a GZV-inducible synTF B (teal), and a GFP reporter unit (green). As in Fig. 2A, 4-OHT (input A) binds to the ERT2 domain (light blue), allowing synTF A to translocate into the nucleus and activate expression of the GFP (output). GZV (input B) inhibits the proteolytic activity of NS3 (orange), preventing the fragmentation of the synTF B and allowing it to GFP expression (output). Right: the multi-input circuit architecture enables cells to perform Boolean logical operations, including OR (top) and AND-logic (bottom). (**B**) The genetic components that comprise the multi-input design space, organized into synTF parts (top), coding parts (middle left), and reporter parts (middle right). Schematic of the assembly workflow, from protein domains to full-length gene circuits (bottom), where the numbers inside each plasmid or plasmid pool represent the number of variants included in the base/full design space. (**C**) Coverage, balance, and indexing for multi-input circuit libraries. Left: circuit (light gray text, bottom) and barcode (dark gray text, top) recovery at each step of the CLASSIC workflow for the base (light blue boxes) and expansion (light gray boxes) libraries. Top right: Nanopore sequencing data of the library to assess composition counts, revealing 7% coverage in the base (light blue histogram) and 0.002% coverage of the expansion (light gray histogram) design spaces. Bottom right: Nanopore sequencing data plotted in rank order by reads per composition (base library: light blue, expansion library: light gray) and barcodes per composition (base library: dark blue, expansion library: dark gray). (**D**) Circuit library sorting and measurement. Top left: GFP expression histograms for base (dotted line) and expansion (shaded) circuit libraries in the presence of no inducer (gray, top left), 4-OHT only (navy, top right), GZV only (orange, bottom left), or both inducers (green, bottom right), overlaid with the boundaries of flow sorting binds (vertical grey dotted lines) and empty HEK293T-LP cells (light gray histogram). Top right: CLASSIC versus ground-truth GFP expression measurements of randomly isolated compositions from the base (n = 14, left) and expansion (n = 13, right) design spaces for all four inducer conditions (no inducer: gray, 4-OHT only: navy, GZV-only: orange, both inducers: green). Bottom: Flow cytometry histograms (top) and CLASSIC-computed distributions (bottom, kernel density (solid line) calculated from normalized barcode read count (shaded vertical bars) for one composition from the base library (dark gray fill in scatter) are shown alongside the accompanying composition ID.

To create a multi-input circuit design space mapping, we first trained a 5-layer MLP (**Fig. 5A**) with the base library data, yielding a model with strong predictive accuracy (90:10 train/validation split with test set *r^2^* values of 0.80/0.90/0.86/0.88 for basal/4-OHT/GZV/both inducer predictions). We then extended the model to the full 3.4B-member design space by fine-tuning with expansion library data, resulting in good predictive performance (*r^2^* values of 0.79/0.84/0.84/0.82) (**Fig. 5B, left**). We calculated similarity of circuit composition behavior across the design space to that of idealized AND- and OR-gates using Kullback-Leibler divergence (see **Methods**). We found that, while only a small region of base library space demonstrated Boolean-like behavior (*D_KL_* values of <0.12 for AND, <0.05 for OR), these regions expanded in the complete design space (**Fig. 5B, right**). To validate the model predictions, we constructed 36 individual compositions spanning the behavior space, 35 of which were unmeasured, including many predicted to exhibit AND- and OR-like function (**Fig. 5C**). As before, we observed good agreement between measurements of individual compositions and the model (*MAE* = 0.32) (**Fig. 5C**), confirming the effectiveness of our data sampling and training approaches.

**Figure 5.**
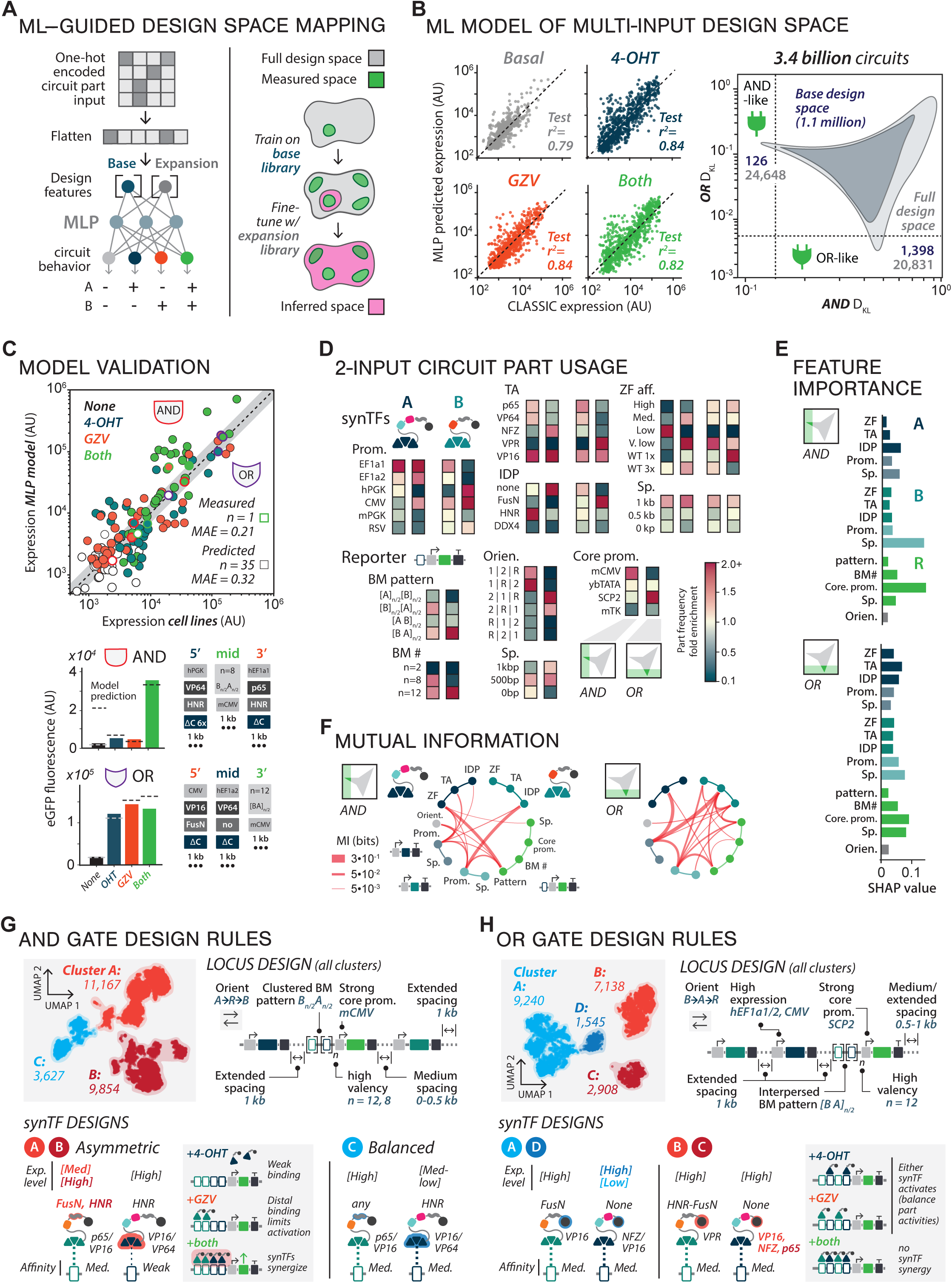
Analysis of digital logic gene circuit design rules in >10^9^ member design space. (**A**) Left: ML model for predicting multi-input circuit behaviors (basal: light gray, navy: 4-OHT only, orange: GZV only, green: both inducers) using OHE part compositions from base (navy) and expansion (dark gray) library data (left). Right: the architecture of the MLP enables exploration of large design spaces (light gray) by first training on a base library (green, top) to infer a small section of the design space (pink, middle) and then fine-tuning using additional data (green, middle) to infer the entire design space (pink, bottom). (**B**) Left: scatter plots of the test set *r^2^* from the fine-tuned model for each of the four input conditions (basal: light gray, navy: OHT only, orange: GZV only, green: both inducers). Right: compositions from the base (dark gray) and full (light gray) design spaces are plotted as Kullback-Leibler divergences (*D_KL_*), which describe the similarity between the model-computed output behaviors and the archetypal AND and OR behaviors. Compositions to the left of the vertical dotted line or below the horizontal dotted line represent the most AND- or OR-like designs, respectively. (**C**) Scatter plot of predicted versus ground-truth GFP expression measurements for 36 individually constructed compositions from across the behavior space (white dots: basal, navy dots: 4-OHT only, orange dots: GZV only, green dots: both inducers) (*MAE = 0.32*) (top). Only 1 of the 36 compositions was measured by CLASSIC. The most AND-like (thick red outline) and OR-like (thick purple outline) designs are highlighted in the scatter plot (top) and broken out below (middle and bottom, respectively). (**D**) Genetic part usage in the AND- (left) and OR-like (right) regions of design space, grouped according to the synTF (synTF A: left, synTF B: right). Part fold enrichment is calculated by dividing the observed part occurrence by expected part occurrence from a balance library. (**E**) Feature importance for all part categories based on absolute SHAP values computed in either AND-like (top) or OR-like (bottom) regions. Larger SHAP values indicate greater contribution to the model’s predictions of AND- or OR-like behavior (navy: synTF A, dark gray: synTF coding, teal: synTF B, light blue: synTF B coding, green: reporter). (**F**) Mutual information between part categories in the AND- (left) and OR-like (right) regions of behavior space, as indicated by red line thickness (navy: synTF A, dark gray: synTF coding, teal: synTF B, light blue: synTF B coding, green: reporter). (**G**) Clustering analysis of the AND-like circuit designs. Top left: UMAP projection of AND-like designs (light red: Cluster A, dark red: cluster B, blue: cluster C). Top right: schematic of the most common locus design observed in AND-like compositions (navy: synTF A, green: reporter, teal: synTF B). Bottom left: schematics of the part usage in synTF B (left) and synTF A (middle) in clusters A and B (light red: Cluster A, dark red: cluster B) and an overview of the dominant strategy for achieving AND-like behavior (right). Bottom right: schematics of the part usage in synTF B (left) and synTF A (right) in the minority cluster (blue: cluster C). (**H**) Clustering analysis of the OR-like circuit designs. (Top left) UMAP projection of AND-like designs (blue: Cluster A, light red: Cluster B, dark red: cluster C, dark blue: cluster D). Top right: schematic of the most common locus design observed in AND- like compositions (navy: synTF A, green: reporter, teal: synTF B). Bottom: schematics of the part usage in synTF B and synTF A (left, middle) in all clusters and an overview of the strategy for achieving OR-like behavior (right) (blue: Cluster A, light red: Cluster B, dark red: cluster C, dark blue: cluster D).

Similar to the single-input design space, we observed strong polarization in part usage within both AND- and OR-like regions (**Fig. 5D**). While compositions from both regions were enriched for a high number of BMs (n=8 or 12) and strong core promoters (mCMV or SCP2), AND-like compositions predominantly utilized clustered BM patterns, while OR gates used exclusively interspersed patterns. In both regions, the 4-OHT-inducible synTF (synTF A) was expressed at intermediate levels (hEF1a1) while the GZV-inducible synTF (synTF B) showed stronger expression and greater overall part diversity. SHAP analysis revealed that core promoter and inter-EU spacing were common drivers of both behaviors (**Fig. 5E**), and OR-like behavior was impacted by more features than AND-like behavior, indicating a greater degree of feature interactivity. This was consistent with part usage coupling analysis, which showed a lower degree of interdependence amongst AND-like compositions, with strong coupling between BM pattern, both synTF expression promoters, and synTF A affinity (**Fig. 5F, left**). The higher inter-category coupling in the OR-like region was driven by strong interactions between features on different synTFs (promoters and TAs) (**Fig. 5F, right**), suggesting co-optimization of expression level and activation strength are important for tuning OR-like circuit behavior.

Clustering analysis of AND-like compositions yielded 3 major clusters. As with HFC circuits, locus design remained constant across clusters (**Fig. 5G**). The largest subset of AND-like compositions (clusters A and B) featured a low-affinity synTF A binding to BMs near the promoter, while synTF B with a medium-affinity ZF and a weak TA bound distally. This suggests a mechanism in which synTF B, separated from the core promoter by ∼110 bp, is unable to independently activate transcription but can facilitate activation by reinforcing weakly-binding synTF A, potentially through IDR-mediated interactions (HNR on synTF A, and FusN [cluster A] or HNR [cluster B] on synTF B) (**Fig. 5G, right**). While this “asymmetric” activation mechanism comprises the majority of AND-like composition, we observed a third, minority solution family (14%) that utilizes balanced synTF expression levels and affinities to achieve AND-like behavior. Clustering of OR-like compositions revealed 4 clusters containing designs allowing synTFs A and B to independently activate expression. This is likely facilitated by interspersed BM binding arrays, which provide both synTFs with similar access to the core promoter (**Fig. 5H**). Medium-affinity synTF binding is a common feature of all 4 clusters, and a lack of IDRs for synTF A suggesting that synTF interaction plays a limited role in reporter activation. In summary, our mapping of multi-input circuit design space allowed for identification of non-intuitive compositional rules, and offered insights into the role that different genetic parts play in determining circuit function.

Here, we have established the utility of combining long- and short-read NGS modalities to perform massively parallel quantitative profiling of multi-kb length-scale genetic part compositions in human cells. CLASSIC holds potential for exploring the emergence of phenotypic function from genetic module composition of across a range of organizational scales and phylogenetic contexts, including for viruses^63^, bacterial operons^64^, and chromatin domains^64–66^. Additionally, the flexibility of our assembly workflow makes CLASSIC adaptable to a wide array of optimization tasks and phenotypic screening strategies. As we show in this piece, by enabling HT profiling of diverse genetic part compositions, our approach significantly expands the scope of inquiry for synthetic biology projects, potentially minimizing the number of DBTL cycles and cost required to identify behavior-optimized part compositions for projects that involve exploration of medium- to largescale design spaces. We envision that CLASSIC could be useful for particularly challenging design tasks in underdetermined systems, wherein iterative rounds of design space expansion can enable convergence to difficult-to-access target behaviors. Though, for smaller-scale projects (<100 constructs) that can leverage well-defined design rules, traditional approaches may be more cost-effective. Furthermore, since CLASSIC leverages extant, broadly-accessible molecular cloning and experimental analysis pipelines, we anticipate it will increase the pace and scale of genetic design for diverse synthetic biology applications across a spectrum of organismal hosts, including the development of bioproduction strains^67^ and multigenic cell therapy programs^5^, as well as other applications that require probing complex genetic design spaces to achieve multidimensional phenotypic optima.

Finally, we showed that data acquired using CLASSIC can not only train ML models to accurately make predictions for out-of-sample and edge-case circuit behaviors but, with properly crafted training strategies, to generalize to orders-of-magnitude larger genetic design spaces. Our results demonstrate the utility of using a data-driven approach to solve complex circuit design challenges, showcasing the ability to identify designs that support target circuit behavior, while also revealing underlying design principles from across design space that are non-intuitive and challenging to capture using biophysical modeling alone. While extensive recent work has leveraged ML approaches to develop sequence-to-function models for various classes of genetic parts^28,68–70^, our work serves as a starting point for developing AI-based models of gene circuit function that use part compositions as learned features. In the future, multi-modal integration of CLASSIC data with sequence-to-function data could be used to train high-capacity deep-learning architectures (e.g., transformers) to enable the generative design of circuits to user-specific function. Such approaches could work in a black-box fashion, without the incorporation of regulatory or biophysical priors, or synergistically with existing mechanistic frameworks to create interpretable models that provide deeper insights into genetic design^71^.

## METHODS

### Plasmid library construction

Circuit libraries were cloned using a custom hierarchical golden gate^35^ assembly scheme, which enables rapid, modular cloning of complex gene circuits starting from individual genetic part sequences. Briefly, input DNA fragments are amplified from genomic or commercial DNA sources and cloned into kanamycin resistant (KanR) sub-part entry vectors using BpiI (Thermofisher) (*level 0*). Sequence-verified sub-part plasmids are then used as inputs for assembly into carbenicillin-resistant (AmpR) entry vectors using BsaI-v2 HF (NEB) to yield genetic part plasmids (i.e., promoters, ORFs, terminators) (*level 1*). EUs are then constructed by assembling promoter, ORF, and terminator part plasmids into a kanamycin-resistant entry vector using Esp3I (Thermofisher) (*level 2*). EUs are subsequently combined into multi-unit arrays by assembling into a carbenicillin – or spectinomycin-resistant (SpectR) destination vectors using BpiI (Thermofisher) (*level 3*).

In this system, barcode pools are incorporated into library assemblies at level 2 and 3. Level 2 barcoded EU pools were generated by first constructing a destination vector carrying a ccdB placeholder expression cassette downstream of the BFP stop codon. A semi-degenerate 18 bp barcode oligo pool (IDT)^72^ was polymerase-extended and cloned in place of the ccdB cassette using BsaI-v2 (NEB) in a 20 µL golden gate reaction: 2 min at 37 °C and 5 min at 16 °C for a number of cycles equal to 10x the number of input fragments, followed by a 30 min digestion step at 37 °C and subsequent sequential 15 min denaturation steps at 65 °C and 80 °C. Resulting assemblies were purified using a miniprep column (Epoch Life Sciences), electroporated (BioRad) into 100 µL NEB 10-Beta electrocompetent *E. coli* (NEB), and plated onto a custom 30 in x 24 in LB_Kan_ agar plate, yielding ∼120M colonies. Plates were grown at 37 °C for 16 h, at which time colonies were scraped into LB_Kan_ (20 mL), incubated for 30 min at 37 °C, and the plasmid library extracted via miniprep (Qiagen). For large-scale libraries, level 3 barcode pools were generated in order to ensure a unique barcode-to-circuit mapping. To create level 3 barcode pools, a semi-degenerate 18 bp oligo pool (IDT) was polymerase-extended and cloned via golden gate assembly into a SpectR destination vector carrying a placeholder ccdB cassette using PaqCI (NEB) following the assembly protocol above. Resulting assemblies were transformed into 100 µL inhouse generated chemically competent^73,74^ Stbl3 *E. coli* cells and plated onto a 10 cm LB_Spect_ agar plate. Plates were grown at 37 °C for 20 h to yield approximately 15,000 colonies, followed by colony scraping and plasmid DNA extraction, as described above.

Part pools (levels 1), EU pools (level 2) and circuit libraries (level 3) were constructed by combining input plasmids at 50 fmol for each part category (15 µL total volume) using the above-described cycling protocol. Transformations varied by library size: level 2 and 3 assemblies for the EU library were respectively transformed into 100 µL or 200µL of chemically competent *E. coli* (DH5a and Stbl3) and plated on LB_Kan_ or LB_Carb_ agar plates for 12-16 h (37 °C) to yield ∼15,000 colonies, respectively. Colonies were scraped into 13 mL LB_Kan_ or LB_Carb_ and miniprepped using a Qiagen kit. Level 1 and 2 assemblies for the single-input library (**Fig. 2B**) were transformed using 100 µL and 400 µL of chemically competent *E. coli* (Stbl3) and grown on LB_Carb_ and LB_Kan_ agar plates at 37 °C for 12-16 h to yield ∼5,000 and ∼20,000 colonies, respectively. Level 3 single-input base library assemblies were purified using a miniprep column (Epoch Life Sciences), electroporated (BioRad) into 100 µL 10-beta electrocompetent cells (NEB), and then plated on 10 15 cm LB_Spect_ agar plates for 16-20 h (37 °C) to yield ∼3M colonies. Each level 3 single-input fine-tuning library assembly was transformed into 500 µL of chemically competent *E. coli* (Stbl3) and grown on LB_Spect_ agar plates at 37 °C for 12-16 h to yield ∼40,000 colonies.

Level 1 assemblies for the multi-input base library (**Fig. 4B**) were treated identically to those from the single-input library. All level 2 assemblies associated with the base design space were transformed into 100µL of chemically competent *E. coli* (DH5a) and grown on LB_Kan_ agar plates at 37 °C for 12-16 h to yield ∼200 colonies. 20 (reporter EUs) or 32 (synTF A or B EUs) colonies from each level 2 library were picked separately into 1 mL of TB_Kan_ and shaken overnight at 37 °C and 900 rpm. 200µL from the 20 or 32 cultures associated with each level 2 library was combined and miniprepped using a Qiagen kit. Separate level 3 base library assemblies were carried out for each of the 6 possible circuit orientations, and purified using a miniprep column (Epoch Life Sciences), electroporated (BioRad) into 15 µL of electrocompetent cells (10-beta, NEB), and then plated on a 15 cm LB_Spect_ agar plates for 16-20 h (37 °C) to yield ∼200k colonies. Cells were scraped and miniprepped separately using a Qiagen kit.

Level 1 assemblies for the multi-input expansion library were generated separately for each new part (24 in total) by combining balanced pools of level 0 part plasmids while holding the new parts constant (**Fig. 4B**). After transforming, scraping, and miniprepping these new part libraries as described above, Level 2 assemblies were generated for each new part, transformed into 20 µL of chemically competent *E. coli* (DH5a), and grown on LB_Kan_ agar plates at 37 °C for 12-16 h to yield ∼50-100 colonies. To sample compositional diversity within each new part library, 10 colonies from each plate were picked into 1 mL of TB_Kan_, shaken overnight at 37 °C and 900 rpm, pooled, and miniprepped using the Star Liquid Handler (Hamilton) to generate 72 level 2 new part libraries. Each of these libraries were then assembled with 5-member level 2 pools from the base library that corresponded to positions not containing the EUs harboring the new parts, along with the BFP barcode EU pool. These level 3 assemblies were then pooled based on their orientation, column purified, electroporated (BioRad) into 7.5 µL of 10-beta electrocompetent cells (NEB), and then plated on a 10 cm LB_Spect_ agar plates for 16-20 h (37 °C) to yield ∼100k colonies. The libraries were scraped and miniprepped separately by Qiagen kit.

### Long-read plasmid sequencing

To generate indices linking assembled constructs to their associated DNA barcodes, we used Oxford Nanopore Technology (ONT) long-read sequencing^75,76^. Libraries were prepared for long-read sequencing by digesting 2 µg of a level 3 plasmid library using Esp3I (Thermofisher) and purifying linearized fragments using a 0.5x volume of magnetic beads (Omega Bio-Tek) by volume. Nanopore sequencing adapters were added to the linearized DNA pool using the LSK-112 or LSK-114 ligation sequencing kit (ONT) and sequenced using on a MinION Mk1b device (ONT) equipped with an R10 flow cell (FLO-MIN112) or R10.4.1 flow cell (FLO-MIN114). Base-calling was performed using Guppy or Dorado (ONT, super high accuracy mode) running on a GPU (Nvidia RTX 3090). Composition and barcode assignment was performed using WIMPY (what’s in my pot, y’all), a custom Matlab analysis pipeline composed of six discrete modules that converts basecalled Fastq files into a part composition array and barcode sequence for each Nanopore read. Briefly, WIMPY first uses Fastqall (1) to import and consolidate basecalled fastq files, followed by Bowtiles (2) to index the reads to a constant region in the level 3 plasmid backbone. In this study, the puromycin resistance cassette (PuroR) was designated as the constant region, and after indexing to PuroR, reads were filtered to remove incomplete assemblies and non-library fragments. WIMPY then calls Tilepin (3), which uses localized containment searching (aka tiling) to identify important genetic landmarks, such as ORFs, to define the different sections of the sequence (e.g., reporter array, synthetic transcription factor coding region, or barcoded BFP expression unit). Chophat (4) then truncates the reads into smaller arrays based on the previously identified landmarks to aid later identification of individual parts. WIMPY then calls Viscount (5a) or FASTar (5b). Viscount determines compositions for filtered reads by identifying and assigning genetic parts through a combination of general Smith-Waterman alignments^77^ and localized containment searches. This is done by splitting the reference sequence into 6-10bp “tiles” with a 1bp stride, after which nanopore reads are queried for the number of tiles contained for each part reference sequence. FASTar uses the same localized containment searches but indexes the location and number of observed reference sequences, and is used to identify repeated sequences, such as number of BMs in the single-input library, or BM patterns in the dual-input library. Reads containing >3% of tiles for a reference sequence are assigned while reads with more than one reference assignment are discarded. Barcode sequences for each read are concurrently determined by Barcoat (5c), in which the sequence section downstream of the BFP EU is aligned to a degenerate reference sequence using a custom alignment matrix.

### Cell culture

HEK293T cells (ATCC^®^ CRL-11268™) used in this study were cultured under humidity control at 37 °C with 5% CO_2_ in media containing Dulbecco’s modified Eagle medium (DMEM) with high glucose (Gibco, 12100061) supplemented with 10% Fetal Bovine Serum (FBS; GeminiBio, 900-108), 50 units/ml penicillin, 50 µg/ml streptomycin (Pen Strep; Gibco, 15070063), 2 mM L-Alanyl-L-Glutamine (Caisson labs, GLL02), referred to hereafter as complete DMEM. HEK293T-LP cells with and without integrated libraries were maintained in complete DMEM supplemented with 50 µg/mL Hygromycin B (Sigma, H3274) and 1 µg/mL Puromycin (Sigma, P8833), respectively.

### Single-copy library integration

To establish a landing pad (LP) cell line, low passage HEK293T cells were co-transfected with a dual Cas9 and gRNA expression vector (pX330-T2) targeting the human *AAVS1* locus^78^ and a PmeI-linearized (NEB) repair template comprising a constitutively expressed YFP-HygR-expression cassette with a BxB1 *attP* recognition site flanked by 800bp homology arms for the human *AAVS1* locus (pROC079). Genomic integration events were selected using 50 µg/mL Hygromycin B, and YFP+ cells were flow sorted to isolate clones (WOLF, NanoCellect). Clones were expanded to 24-well plates and tested for integration competency by integration of an *attB*-containing destination vector and subsequent flow cytometry and the presence of the intact LP cassette at the *AAVS1* locus by PCR. gDNA was extracted from each clonal population (Omega Bio-Tek) and 50 ng of gDNA was used as the template in a 25 µL reaction along with 0.25 µL of Q5 polymerase (NEB), 0.5 µL of 10mM dNTPs, 5 µL of 5X Q5 polymerase buffer, and 1 µL of each 1 mM genotyping. Primers were specific to the native AAVS1 locus, Landing Pad, or destination vector. The thermocycling protocol was 2 min at 98 °C, then 30 cycles of 98 °C for 20 s, 65 °C for 20 s, 72 °C for 1 min and then a final step of 72 °C for 5 min. We identified a clone (LP79 #36) that demonstrated BxB1-dependent, single copy integration of vectors harboring an *attB* recognition site and used this cell line for all following experiments. We verified single-copy genomic integration of the landing pad cassette by ddPCR using primer sets for the engineered locus (Hygro) and a pair of previously validated native genes (Beta Actin and POTP). Briefly, each 25 µL reaction was composed of 12.5µL of EvaGreen Supermix (BioRad), 1.25µL of each primer (diluted to 2µM), 20ng of gDNA, and 0.1 µL of HindIII (NEB). Droplets were generated in the QX600 Droplet Generator (BioRad) by combining 20 µL of each reaction with 70 µL of Droplet Generation Oil (BioRad). PCR was performed on 40 µL of the emulsified reaction using the following protocol: 5 min at 95 °C, then 40 cycles of 95 °C for 30 s and 60 °C for 1 min, followed by 5 min at 4 °C and 5 min at 90 °C. The PCR results were then analyzed using the QX600 Droplet Reader (BioRad). All resulting concentrations were normalized to the HEK293T Beta Actin control.

To integrate individual constructs, 1.25 × 10^5^ cells were plated into a 24-well plate one day prior to transfection. The well was co-transfected with 250 ng of a level 3 vector containing a construct of interest and 125 ng of BxB1 expression plasmid (Addgene #51271) using JetPrime (VWR). pCAG-NLS-HA-Bxb1 was a gift from Pawel Pelczar (Addgene plasmid #51271)^79^. Two days after transfection, cells were passaged into complete DMEM containing 1 µg/mL Puromycin and selected for 10 days to achieve homogenous expression. Library integration was performed by co-transfecting HEK293T-LP with 250 ng of plasmid library and 125 ng of BxB1 per 200k cells (384-member library = 1M cells transfected; 166k-member base library = 100M cells transfected; 166k-member sub-library = 10 M cells transfected; 3.4B-member base library = 200 M cells transfected; 3.4B-member expansion library = 200 M cells transfected) using JetPrime. Media exchange was performed after 8 h and cells were expanded for an additional 40 h, split into five culture dishes, and grown for 6-8 days under Puromycin selection. Cells were passaged 1:3 and cultured for an additional 5 days under puromycin selection, and then combined in complete DMEM for flow sorting (10M cells/mL).

To determine the clonal variability within LP-integrated cell populations, we integrated a constitutively expressed mCherry EU following the protocol described above. We then randomly sorted 23 clones from the population (WOLF, NanoCellect) and measured their mCherry expression. We calculated the MAE for this set (*MAE = 0.08*) and used this value to define an expected error range from clonal heterogeneity (ERCH), which represents the expected variability of cells from an integrated population. For the 384-member library, the expression ERCH was set to 0.08 on either side of the expected value. For all libraries constructed as part of the single-inducer circuit class, the fold-change ERCH was calculated to be 0.16 on either side of the expected value. For libraries associated with the multi-input design space, the expression level ERCH was calculated to be 0.16 on either side of the expected value.

### Flow sorting

To prepare libraries for flow sorting, cells were lifted using TrypLE (Gibco), washed twice with PBS, and resuspended in complete DMEM (10M cells/mL). For the 384-member library, the mRuby expression distribution was divided into 10 evenly log-spaced bins on a Sony MA900 set to purity mode. The number of cells collected for each bin was proportional to the percentage of the mRuby distribution expressed in that bin, with a target of ∼125k cells sorted in the most populous bin [e.g., 2,252 cells were collected for bin 1 (0.53% of library), 126,562 for bin 4 (29.78% of library)]. For the single-input base and fine-tuning libraries, cells were split and either grown in DMEM containing 1µM 4-OHT (Sigma Aldrich) or without 4-OHT for 72 h. The eGFP expression distributions for both conditions were then divided into 8 evenly log-spaced bins. Cells were sorted into bins in proportion to the relative abundance of the library in each bin, with the most populous bin set to collect 3 M cells and ∼0.5 M cells in the single-input and fine-tuning libraries (**Fig. 2D**), respectively. For the multi-input base and expansion libraries, cells were split and either grown in DMEM containing 1µM 4-OHT (Sigma Aldrich), 20µM GZV (MedChemExpress), 1µM 4-OHT and 20µM GZV or without inducers for 72 h. eGFP expression distributions for each condition were then divided into 8 evenly log-spaced bins and sorted in proportion to abundance, with the most populous bin set to collect ∼1.5 M cells and ∼1 M cells for the base and expansion libraries, respectively. For all libraries, the sorted cells were plated into 96-, 24- or 6-well plates to achieve a plating density of 10-30%, grown under puromycin selection for 2 d, passaged and washed with PBS, and grown for an additional 3 d. Cells were lifted using TrypLE and total RNA was extracted using the Takara RNA plus kit following manufacturer’s instructions (Takara). 5 µL of total mRNA was converted to total cDNA using random hexamers following manufacturer’s instructions (Verso, Life Technologies).

### Short-read (Illumina) sequencing

To prepare the sorted libraries for short-read sequencing, barcode regions were PCR amplified from cDNA in a 15 µL reaction with Phantamax polymerase (Vazyme) and custom bin-specific primer sets (IDT) using the following thermocycling protocol: 2 min at 98 °C, then 12-18 cycles (depending on the library) of 98 °C for 20 s, 60 °C for 10 s, 66 °C for 10 s, 72 °C for 20 s and then a final step of 72 °C for 2 min. The resulting amplicon pool was extracted from a 2% agarose gel. A second PCR step, which used 6-10 cycles (library dependent) of the thermocycling protocol for PCR step 1, added Illumina sequencing adapters (i5 and i7) and sample-specific sequencing barcodes using Phantamax and custom primer sets (IDT), and the amplified product was extracted from a 2% agarose gel. Purified amplicons from each bin were pooled at equimolar concentrations and sequenced using an Illumina MiSeq (kit v2, 300 cycles), NovaSeq6000 (Sp v1.5 kit, 300 cycles) and the NextSeq 2000 (P2 kit, 200 cycles) for the 384-, 166k-, and 3.4B-member (both base and expansion) libraries, respectively. Illumina data were analyzed using a custom analysis pipeline (Matlab). Briefly, fastq files are imported and split according to their sample-specific sequencing barcode. Barcode sequencing data is converted to average expression for all compositions/variants as follows: (1) Counts for each barcode in each sample/bin are normalized by the percentage of the library distribution in that bin to account for differences in read depth across bins. (2) A weighted average of number of reads in each bin for each barcode is then calculated, to assign an average expression level (using a bin → expression conversion table) to each barcode. (3) Barcode expression levels are then cross-referenced with the Nanopore indices to assign average expression values for all barcodes associated with a given variant in a n-by-y cell array, where n is the number of library variants and y is the (variable) number of barcodes for each variant. (4) For each variant, a gaussian kernel density estimation is performed over the log-normalized expression values from all barcodes associated with each variant, using the Matlab function *ksdensity*. The expression value corresponding to the kernel peak is assigned as the mean expression value for each variant.

### Experimental validation of CLASSIC measurements

To assess the accuracy of CLASSIC measurements for both the EU and single-input libraries, single cells were sorted from the integrated cell populations into 96-well plates (MA900) and clonally expanded from across the mRuby and eGFP expression distributions, respectively. Isolated clonal populations, which are referred to as ‘isolates’, were expanded to 24-well plates and the fluorescence distribution was measured by flow cytometry (MA900). For the EU library, one 96-well plate of cells was sorted based on mRuby expression. For the single-input base library, two 96-well plates of cells were sorted based on eGFP expression from the uninduced cell library. For the single-input fine-tuning library, one 96-well plate of cells was sorted based on eGFP expression from the uninduced population. For both the multi-input base and expansion libraries, two 96-well plates of cells were sorted based on eGFP expression from the uninduced cell libraries. Following fluorescence measurement under library-specific experimental conditions, gDNA was extracted from each clonal population (Omega Bio-Tek). For the EU and single-input libraries, the genetic variant in each isolate was identified by PCR-based amplification and Nanopore sequencing of the LP locus using a forward primer in puromycin resistance cassette and a reverse primer in the BFP ORF. A set of 30 different forward primers, each with defined barcodes, were paired with a common reverse primer to amplify the genetic variants from each isolate and associate the genetic variant with the isolate during Nanopore sequencing. 50 ng of gDNA was used as the template in a 20 µL reaction along with 10 µL of 2X PhantaMax Master Mix (Vazyme), 1 µL nmol of primer, and brought up to 20 µL in nuclease-free H_2_O. The thermo-cycling protocol was 2 min at 98 °C, then 30 cycles of 98 °C for 20 s, 65 °C for 20 s, 72 °C for 5 min and then a final step of 72 °C for 10 min. Each PCR reaction was extracted from a 1% agarose gel and column purified using QG buffer (Qiagen), following manufacture instructions. Purified amplicons were pooled and prepared for Nanopore sequencing using LSK-112 or LSK-114 ligation sequencing kit (ONT) and sequenced with the R10 (FLO-MIN112) or R10.4.1 (FLO-MIN114) MinION flow cell. Basecalling was performed using Guppy or Dorado (ONT, super high accuracy mode) running on a GPU (Nvidia RTX 3090). Composition and barcode assignment was performed using WIMPY, as described above. This process yielded 15 isolates from the EU library, 40 isolates from the single-input base library (**Fig. 2D, middle**), and 8 isolates from the single-input fine-tuning. Isolates from the base and expansion 3.4B-member libraries, circuit identity was determined by amplification of the barcode sequence from gDNA, following the PCR protocol described above, extraction from a 2% agarose gel, and Sanger. Using the CLASSIC composition-to-barcode IDs to assign barcodes to circuit variants yielded 14 and 13 mapped isolates from the base and expansion libraries, respectively.

### MLP deep neural network regression for circuit behavior prediction

#### Model architecture and parameters

Fully connected multi-layer perceptron (MLP) neural network architectures were constructed for single- and multi-input design spaces. The single-input model had 4 hidden layers with 160/80/40/20 neurons, a 39 x 1 input channel, and 2 output channels for log10 transformed basal and induced expression. The multi-input model consisted of 5 layers 160/80/40/20/10 neurons, a 68 x 1 input channel, and 4 output channels. Input compositions were one-hot encoded for each part category and flattened into a 39 x 1 shape for the single-input and 68 x 1 shape for the multi-input library. Stochastic gradient descent with momentum (sgdm) was used to optimize the root mean squared error loss function during training (momentum parameter=0.9), learning rate of 5 x 10^-2^, and a piecewise learn rate scheduler that drops the learning rate by a factor of 0.5 every 10 epochs. The batch size was 256.

#### Data splits for model training

High-quality test sets were constructed from the single- and multi-input base library datasets (121,292 and 88,551 measured compositions, respectively), by randomly sampling 15% of compositions with >12 and >50 barcode-indexed reads (1,868 compositions and 953 compositions respectively). Flow cytometry measurements for isolates were removed from training datasets to allow for unbiased comparisons against these measurements later. The remaining data was randomly split 90:10 into training:validation sets containing 103,878 compositions and validation sets containing 11,532 compositions for the single-input library, and 48,031 and 5,336, respectively, for multi-input library. The validation set was used to monitor training progress, and for early stopping with a patience parameter of 300 iterations. The model with the best validation performance was returned and used to evaluate performance on the high-quality test and isolate sets.

#### Data downsampling

For demonstrating the single-input model’s tolerance to lower amounts of training data (**Fig. 3F**), compositions were downsampled in increasing order of the number of barcoded reads per composition, retaining compositions from ≥ 1 read to ≥ 40 reads to simulate scenarios in which lower amounts of higher-quality data is collected. The model was re-initialized and trained with a new 90:10 training-validation split from the remaining dataset for each of the 40 downsampling iterations, which were all evaluated using the high quality test set mentioned above.

#### Design space expansion via neural network fine-tuning

To demonstrate single-input model fine-tuning, an MLP model with the same architecture and hyperparameters was trained on the original library, with two extra input encoding slots to represent new parts that would be added during fine-tuning. For the multi-input library, the base model already contained extra input slots for new parts. A high-quality test set was created, as described above, from the expansion library datasets, with an increased barcoded read count threshold of 40 (single-input) or 50 (multi-input). The remaining data was split 90:10 into a training and validation temp sets. These sets were combined with 2,000 compositions from the base training set and 500 compositions from the base validation set to avoid memory loss during training. The combined validation set (with base and new compositions) was used for early stopping and all hyperparameter optimization as described above, except for the initial learning rate, which was changed to 5 x 10^-3^. Model performance was evaluated on the high-quality test sets and used to construct individual variants across the behavior space.

### Mutual information calculation

For a given pair of features in a region of design space, mutual information was computed using the following formula:

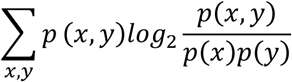

where x and y are 2 different features of the library, and p(x, y) represents the joint frequency for specific x and y combinations divided by the total number of points in the dataset. p(x) and p(y) represent the fractional frequency of the corresponding parts by themselves.

## SHAP & UMAP clustering

For both the single- and multi-input design spaces, SHAP values were used to assess the contribution of each feature in the HFC and AND/OR-like regions of behavior space, respectively. For each model, SHAP values were computed by fitting a SHAP explainer model across 500 randomly sampled training points and then predicting on 200 randomly points sampled from the regions of interest. These values were averaged across expression conditions and within part categories to generate region-specific SHAP scores for each part category.

For the 166k-member library, UMAP dimensional reduction was performed on part composition data from variants in the HFC (>25-fold region) region of the behavior space, using the *run_umap* function in Matlab with a Jaccard distance metric. To identify the optimal number of clusters, a gap test was carried out on the 2D projection using the *clusterevaluation* function in Matlab. HFC compositions were assigned to clusters using the *kmeans* function in Matlab. Cluster stability was determined by performing 100 independent rounds of UMAP projection and k-means clustering, recording cluster affiliation in each iteration, and computing the cluster stability index (CSI) and adjusted rand index (ARI). We then assigned compositions to clusters the most stable UMAP projection from the 100 runs mentioned above. Clustering for AND-like and OR-like circuits from the multi-input library was performed as described above for the single-input library, including the computation of cluster stability. For display purposes, boundaries have been manually drawn around each cluster.

### KL-divergence computation

For multi-input circuits, expression levels were normalized to relative abundances by dividing them by the summed expression across the four induction conditions (no inducer/4-OHT/GZV/both) creating probability distributions suitable for computing Kullback-Leibler divergence (*D_KL_*) against “ideal” AND and OR gates, using the following equation:

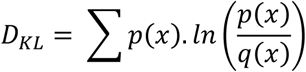

Where *p(x)* represents the normalized distribution across the four conditions, and *q(x)* refers to the probability of the ideal AND [0.01, 0.01, 0.01, 0.97] and OR [0.01, 0.33, 0.33, 0.33] gate across the four induction conditions. Low *D_KL_* values indicate high similarity to ideal AND and OR distributions.

## ACKNOWLEDGEMENTS

We thank Oleg Igoshin, Ankit Patel, Yashwanth Lagisetty, Satpreet Singh, and members of the Bashor lab for helpful discussions. This work was supported by grants from NIH R01 EB029483 (C.J.B.), NIH R01 EB032272 (C.J.B.), ONR N00014-21-1-4006 (C.J.B.), and funding from the Robert J. Kleberg Jr. and Helen C. Kleberg Foundation (C.J.B.). R.W.O. was supported by a graduate fellowship from the American Heart Association (917746). B.K. was supported by a NLM Training Program in Biomedical Informatics and Data Science fellowship (T15LM007093-31) and by NIH grant P01-AI15299901. K.D.C. was supported by NSF EF-2126387 and the Ken Kennedy Institute Computational Science & Engineering Recruiting Fellowship. P.M. and J.W.R were supported by NIH R35GM119461 (P.M.)

## AUTHOR CONTRIBUTIONS

R.W.O., K.R., and C.J.B. conceived of the study, R.W.O. and K.R. carried out the experiments and developed the analysis software, with assistance from T.C.P., Y.W., L.C.B, K.D.S., J.A.W., S.L., T.H.Z., E.M.R., and A.S. R.W.O., and T.C.P. developed the modular cloning scheme and landing pad cell line, with assistance from S.L. B.K., K.C., and T.J.T. helped develop the barcoding scheme and analysis software. R.W.O., K.R., T.C.P., Y.W., J.W.R., P.M, and C.J.B. analyzed the data. C.J.B. supervised the study. R.W.O., K.R., and C.J.B wrote the manuscript, with input from all authors.

## DATA AVAILABILITY

All raw data used in this study is available upon request. Accession codes will be made available prior to publication.

## CODE AVAILABILITY

All custom scripts used for Nanopore sequencing data analysis are available on GitHub (github.com/cbashorlab/WIMPY). Code associated with Illumina data analysis and model training are available upon request.

## CORRESPONDING AUTHOR

Correspondence and requests for materials should be addressed to Caleb J. Bashor.

